# Integration host factor bends and bridges DNA in a multiplicity of binding modes with varying specificity

**DOI:** 10.1101/2020.04.17.047076

**Authors:** Samuel B. Yoshua, George D. Watson, Jamieson A. L. Howard, Victor Velasco-Berrelleza, Mark C. Leake, Agnes Noy

**Affiliations:** Department of Physics, University of York, York YO10 5DD, United Kingdom; Department of Biology, University of York, York YO10 5DD, United Kingdom

## Abstract

Nucleoid-associated proteins (NAPs) are crucial in organizing prokaryotic DNA and regulating genes. Vital to these activities are complex nucleoprotein structures, however, how these form remains unclear. Integration host factor (IHF) is an *Escherichia coli* NAP that creates very sharp bends in DNA at sequences relevant to several functions including transcription and recombination, and is also responsible for general DNA compaction when bound non-specifically. We show that IHF–DNA structural multimodality is more elaborate than previously thought, and provide insights into how this drives mechanical switching towards strongly bent DNA. Using single-molecule atomic force microscopy and atomic molecular dynamics simulations we find three binding modes in roughly equal proportions: “associated” (73° of DNA bend), “half-wrapped” (107°) and “fully-wrapped” (147°), only the latter occurring with sequence specificity. We show IHF bridges two DNA double helices through non-specific recognition that gives IHF a stoichiometry greater than one and enables DNA mesh assembly. We observe that IHF-DNA structural multiplicity is driven through non-specific electrostatic interactions that we anticipate to be a general NAP feature for physical organization of chromosomes.

## Introduction

Nucleoid-associated proteins (NAPs) are a collection of DNA-interacting proteins that perform crucial roles of organization, packaging and gene regulation in prokaryotic chromosomes (1, 2), including functions as transcription factors (3). They often bind non-specifically, depending on their relative concentration in cells, which vary across orders of magnitude depending on cell-cycle stages and environment conditions (4). NAPs are molecular-architectural proteins that can create a wide variety of 3D genomic arrangements (5) by essentially bending and bridging one or more molecule of DNA (6). Although one binding mode (bridging or bending) is usually exclusive of specific recognition for each individual NAP, their entire DNA-interacting activity seems to be far more versatile and promiscuous than previously thought (1). A good illustration of this is with the NAP Fis, which induces bending on DNA when bound in specific sequences (7), but it also can stabilize loop crossings via bridging the DNA (8).

Integration host factor (IHF) is a key NAP in *Escherichia coli* and other Gram-negative bacteria. Its architectural role is thought to involve creating some of the sharpest bends observed in DNA (9), in excess of 160° (10), at around 300 sites containing the consensus sequence WATCARNNNNTTR (W is A or T; R is A or G; N is any nucleotide), thereby facilitating the assembly of higher-order nucleoprotein complexes (11) such as gene regulatory loops (12), the CRISPR-Cas system (13), the origin of replication (*oriC*) (14), and a Holliday junction complex involved in the integration and excision of phage λ DNA (15). IHF’s large repertoire of roles supports the long-standing view that it has an essential function in the structural organization of DNA in a wide variety of genetic transactions. However, with copy numbers on the order of tens of thousands per cell depending on the growth phase (16), non-specific binding must also play a role despite a 1000-fold larger *K_d_* (17), and IHF alone is able to compact DNA (18, 19). This non-specific binding may also play a role *ex vivo*, as IHF has been implicated in biofilm stability of important pathogens like *E. coli* (20), *P. aeruginosa* (21) and *B. cenocepacia* (22). In some cases, the removal of IHF caused a 50% reduction in biofilm thickness (22) and IHF has also been imaged at vertices of an extracellular DNA lattice (21).

The crystal structure of IHF is obtained from its conformation bound to DNA at a specific binding site (10). It revealed that IHF is formed by a core of α helices with a pair of extended β-ribbon arms whose tip each contains a conserved proline that intercalates between two base pairs (Figure 1). These two intercalations stabilize strong bends 9 bp apart and facilitate wrapping of two DNA ‘arms’ around the protein body, tightened by electrostatic interactions between the phosphate backbone and isolated cationic amino acids on the protein’s surface, resulting in a binding site with a total length of 35 bp and an overall bend angle of 160° (see Figure 1), which has been supported by atomic force microscopy (AFM) (23, 24). IHF’s consensus sequence is located on the right side of the binding region and is small compared to the total length of the wrapped DNA. However, most of the strongest IHF binding sites include an A-tract to the left-hand side (upstream of the specific sequence, see Figure 2) that increases affinity, the degree of bending and the length of the attached DNA site (25).

**Figure. 1.**
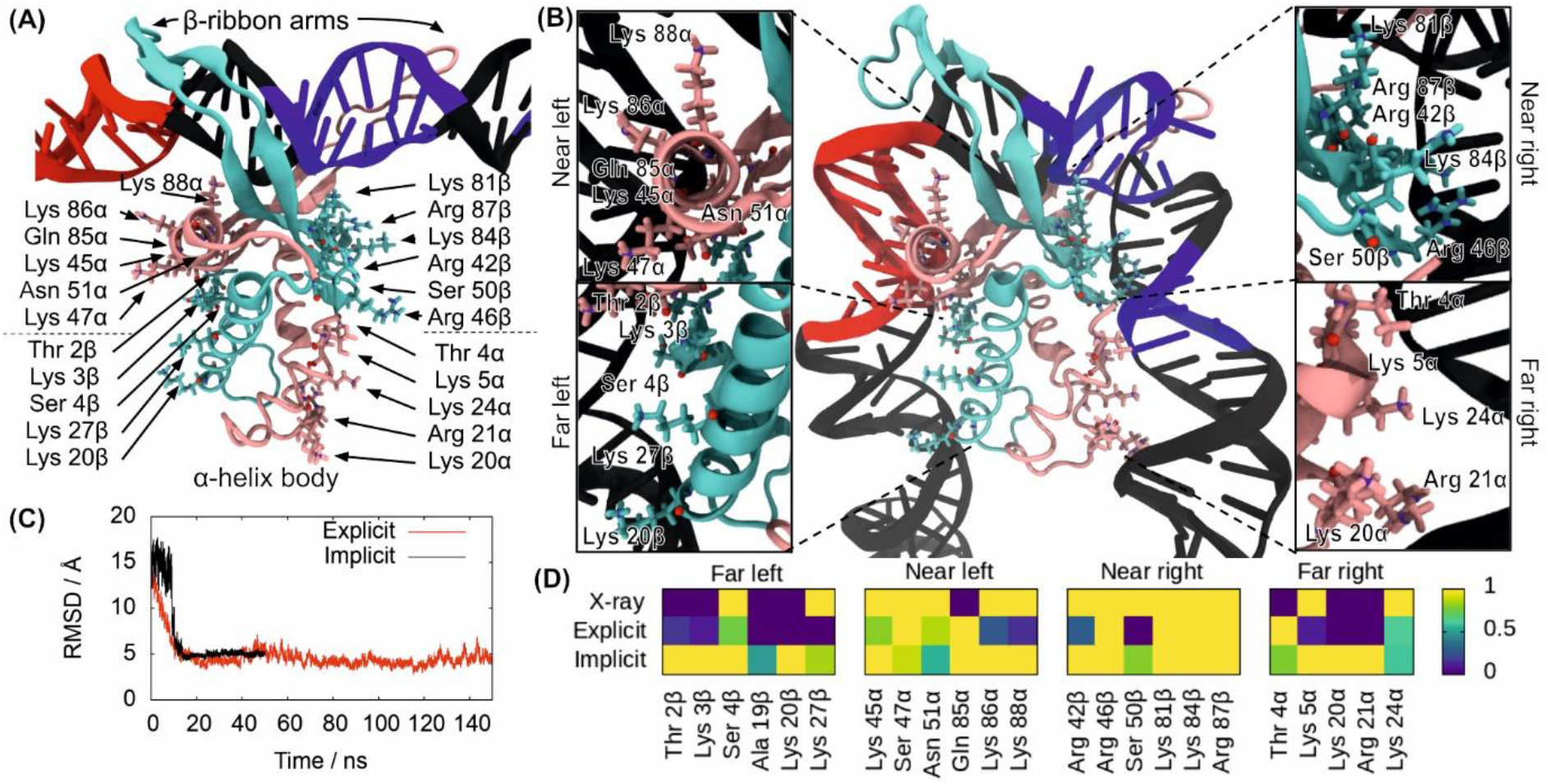
MD simulations of the process of DNA wrapping around IHF. Initial **(A)** and final **(B)** conformations of MD simulations in implicit solvent, with the α subunit shown in mauve and the β subunit in turquoise. The DNA is in black except for the consensus positions (in blue) and A-tract (in red). The amino acids that interact with the lateral DNA arms in the 1IHF crystal structure are labelled and highlighted with atomic representation (N in blue, O in red). **(C)** Time-evolution RMSD of trajectories in implicit and explicit solvent shows transition to a structure close to the PDB 1IHF. **(D)** DNA–protein hydrogen bonds observed in the 1IHF crystal structure are also present in the trajectories on both solvent models. The number of hydrogen bonds is capped at 1, so time-averages along simulations indicate the ratio of frames presenting interaction.

**Figure. 2.**
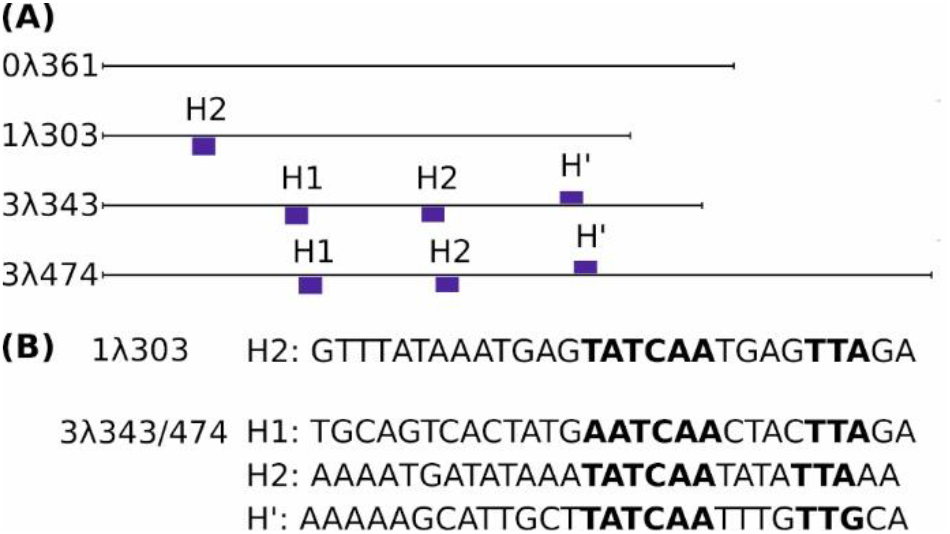
DNA constructs and IHF binding sites used in experiments. Scheme for the different DNA fragments shown to relative scale with IHF binding sites labelled (A) and their sequence specified (B). Only the sites with p-values lower than 0.0002 on the PSSM scanning analysis are displayed by blue bars, which are sized according to the primary score (see Materials and Methods). The most conserved positions of the IHF specific binding sites are in the right-hand side (highlighted in bold) although the presence of A-tracts in the left-hand side make the sequence more affine (25).

Previous work has shown binding occurs through a two-step process, a fast ^~^100 μs step that seems to be not sequence-specific, and a slower millisecond step which is site-specific (26). This second step seems to be associated with an activation energy of approximately 14 kcal/mol that would be related with proline intercalation (27), but the free energy of the wrapping process (Figure 1) is found to be only around 3.6 kcal/mol (28). Recent studies have shown that IHF can bend DNA flexibly with a portion of the population that is only partially bent (i.e. <160°) (29, 30), demonstrating that the complex is more dynamic than originally thought. Despite these advances, the underlying bending mechanism driven by IHF is still poorly understood, with no obvious explanation as to why the apparent free energy of DNA wrapping is so close to the thermal energy scale.

Here, we compared experimental AFM imaging with atomically precise predictions from molecular dynamics (MD) simulations for the same DNA sequences of around 300 bp (Figure 2). This approach allowed us to show how a strong binding site, H2 (17, 31), has a multiplicity of binding modes beyond the canonical 160° state, which is the structure resolved by crystallography. We then formulated a model for multiple IHF bending states encapsulating both specific and non-specific binding. Using advanced modeling methods, we calculate the free energy of each of these conformations and demonstrate asymmetric behavior between DNA arms that allows different mechanical switching depending upon the order of DNA arm binding. Furthermore, by looking at the effects of increasing the number of IHF binding sites using AFM, we see evidence for IHF-mediated bridging between two DNA strands at low protein concentrations and provide structural insight into how this occurs. The variety of DNA binding modes caused by IHF – both bending and bridging - may be representative of the behavior between DNA and other nucleoid-associated proteins.

## Materials and Methods

### Protein amplification and purification

IHF was overexpressed from the *E. coli* strain BL21AI containing the plasmid pRC188 (a kind gift from the Chalmers laboratory, the University of Nottingham, UK). The cells were grown in 2x 1 L LB + 100 μgml^−1^ carbenicillin at 37°C with shaking at 180 rpm to an OD_600_ ^~^ 0.6. Overexpression of IHF was induced by the addition of IPTG and Arabinose to final concentrations of 1 mM and 0.2% (w/v) respectively and growth was allowed to proceed for a further 3 hours. Cells were harvested by centrifugation at 4,000 rpm and 4°C in a Sorvall SLC6000 rotor. The pelleted cells were then resuspended in 20 mL of 10 mM Tris pH 7.5, 10% sucrose (w/v) before being flash frozen in liquid nitrogen and stored at −80°C.

For purification stored cell pellets were thawed on ice and the buffer was adjusted to contain 50 mM Tris pH 8.4, 150 mM KCl, 20 mM EDTA, 10 mM DTT and 0.2 mgml^−1^ lysozyme. The resultant suspension was mixed by inversion and left on ice for 15 mins before the addition of Brij58 to 0.1% (w/v) and a further 15 minutes on ice. The suspension was then clarified by centrifugation at 4°C and 38,000 rpm in a Beckmann Ty70Ti rotor for 60 minutes. Polymin P was added to 0.075% (w/v) from a 1% stock in a dropwise fashion to the supernatant whilst stirring at 4°C, stirring was continued for 10 mins before centrifugation at 4°C and 16,000 rpm in a Sorvall SS34 rotor for 20 minutes. The supernatant was collected before being subjected to a 50% ammonium sulfate (AmSO4_4_) precipitation followed by an 80% AmSO4_4_ precipitation. In each case the sample was centrifuged as above IHF remained soluble at 50% AmSO4_4_ and precipitated at 80% AmSO4_4_. The precipitated IHF was resuspended in 20 ml buffer A (50 mM Tris·HCl pH 7.5, 2 mM EDTA, 10 mM β-ME, 10% glycerol) such that the conductivity matched that of buffer A + 100 mM KCl. The sample was loaded onto a 10 mL P-11 phosphocellulose column equilibrated with buffer A + 100 mM KCl, the column was washed with 300 ml of buffer A + 100 mM KCl before being developed with a 200 mL gradient of 0.1-1 M KCl in buffer A. Fractions containing IHF were identified using 15% SDS polyacrylamide gel electrophoresis and pooled. The pooled fractions were dialyzed against buffer A + 100 mM NaCl before loading onto a 5 mL Hitrap Heparin column (GE Healthcare) equilibrated with the same buffer, the column was washed with 300 ml of buffer A + 100 mM NaCl before being developed with a 200 mL gradient of 0.1-1 M NaCl in buffer A. Fractions containing IHF were again identified using 15% SDS polyacrylamide gel electrophoresis and pooled. Pooled fractions were aliquoted and flash frozen in liquid nitrogen before storage at −80°C. Protein concentrations were determined using the Bradford Protein Assay (Bio-Rad).

### Production and composition of DNA fragments

DNA constructs were chosen from phage λ with different numbers of IHF binding sites, within the range of 300-400 base pairs. All DNA constructs were amplified by PCR using Q5 DNA polymerase (see Table S1 for primers) resulting in three constructs: 0λ361 (361 bp long containing no IHF consensus sequence), 1λ306 (306 bp long with one IHF consensus sequence) and 3λ343/478 (343/478 bp long with three IHF consensus sequences). These were all chosen in the region around the *xis* gene that is known to interact with IHF (32). The two constructs with three binding sites are highly homologous presenting a perfect overlap of 3λ343 sequence to 11-351 position on 3λ478 (Figure S1).

The presence of sites with significant similarity to the IHF consensus sequence was evaluated by scanning the position-specific scoring matrix (PSSM) available at the Regulon database (33) along our constructs using the program *matrix-scan* from the regulatory sequence analysis tools (RSAT) web server (http://rsat.ulb.ac.be/rsat/) (34). The algorithm assigns a score (W_s_) to all possible segments (S) defined by W_s_=log_2_(P(M)/P(B)), which evaluates the ratio of probabilities to find a particular sequence in our PSSM (P(M)) with respect to a sequence background (P(B)) (the whole *E. coli* genome in our case) (34). The program also calculates the strength of a match using a p-value, which evaluates the risk of false positives. We recovered the known specific binding sites with a p-value cut-off of 0.0002 (see Figure 2). A more permissive scanning, which considered all matches with W_s_≥1 or P(M)≥2P(B), revealed additional secondary sites, although practically none of them were in 0λ361 (see Figure S1). This demonstrates the absence of any binding site with a resemblance to the IHF consensus sequences in this construct.

### Atomic force microscopy acquisition and analysis

Mica was freshly cleaved and then pre-treated by depositing 20 μl 0.01% (w/v) 1-5 kDa poly-L-lysine (Sigma-Aldrich, MO, USA), left for 5 mins before washing with 1000 μl filtered milliQ H_2_O and finally vacuum drying. The DNA samples were prepared by adding a 20 μl solution of 10 mM Tris, 50 mM KCl with 1 nM DNA (and 5 – 150 nM IHF) to the pre-treated mica and leaving for approximately 5 mins. These were then washed once more with 1000 μl of filtered milliQ H2O and vacuum dried before being imaged.

All samples were imaged on a Bruker Bioscope Resolve (Bruker, CA, USA) in tapping mode using TAP-300AI-G probes (BudgetSensors, Bulgaria). Images were taken at 512×512px, with a pixel size of 3.9 nm. These were then loaded using pySPM (35) and preprocessed by using a 1st order line-by-line flattening and 2nd order fits to flatten the surface before filtering scars (36).

To characterize individual DNA strands, images were then analyzed using custom code using the scikit-image package (37) to threshold the image. The individual segments were then skeletonized to recover the DNA contour for further analysis. Segments whose length were not within 50% of the anticipated contour length (such as debris, aggregates, “spot-like” DNA or globular structures caused by the deposition of poly-L-lysine in mica (38)) were discarded for the subsequent analysis. They accounted for approximately 15% of the total. This heuristic threshold was sufficient to remove outlier objects (too short or too long) but did not result in any significant deletion of non-aggregated DNA molecules as indicated in Figure S2. The average contour length per base pair was found to be around 0.30 nm (10% smaller compared with the nominal B-DNA value of 0.34 nm), which is in agreement with the previous identified value for DNA absorbed on mica (39). The number of DNA molecules analyzed were *n=87/63 for 0*λ361 +/- IHF and *n*=144/584 for 1λ306 +/- IHF. The initial AFM experiments using the 0λ361 construct required front-loading of lab access time for significant optimisation to get high quality imaging data, whereas later datasets could then use these optimised imaging and incubation conditions enabling greater numbers of high quality data to be obtained.

One angle was measured per each individual DNA strand, first identifying the largest peak along the DNA contour, which was IHF if present or random if not, and then taking a 4 px (16 nm) vector either side of a 3 px window around the peak. This heuristic method was designed to account for the limited resolution in our AFM setup, partially caused by the similar effective diameter of IHF compared to the width of the DNA. Once angular distributions were determined, goodness-of-fit tests were carried out, with *p* > 0.99, where *n=87/63 for 0*λ361 +/- IHF and *n*=584/144 for 1λ306 +/- IHF (see Figure S3 for more information). Because molecules were kinetically trapped in the 2D instead of being equilibrated (40), bend angles from AFM could only be related to the 3D structure through the comparison with simulations (see below).

For the analysis of clusters/aggregates the zero basis volume (see Gwyddion documentation (41)) of each identified segment (before skeletonization) was calculated, filtering any cluster smaller than 6 pixels.

### Unbiased molecular dynamics simulations

All simulations were set up with the AMBER18 suite of programs and performed using the CUDA implementation of AMBER’s pmemd program (42). A linear B-DNA molecule with a sequence of length 302 bp extracted from the *xis* gene of bacteriophage λ was generated using the NAB utility. This sequence is highly homologous to the 306 bp experimental construct obtained using commercially available λ DNA (New England Biolabs), so both are herein referred to as 1λ306 to minimise confusion. The both sequences have 100% sequence identity apart from four additional base pairs close to the ends, which represents less than 2% of the total. IHF bound to the H2 binding site was extracted from PDB entry 5J0N (15). Only the eleven base pairs at the center of the binding site were maintained with the purpose of starting the simulations with unbent DNA just attached to the extended β-ribbon arms of IHF (Figure 1A). The complex was then inserted at the relevant location of the linear DNA molecule. A 61 bp section of the 1λ306+IHF construct, centered on the binding site, was explicitly solvated using a truncated octahedral TIP3P box and neutralized with a 0.2M-equivalent concentration of K and Cl ions (43). The protein and DNA were represented using the ff14SB (44) and BSC1 (45) force fields, respectively. Simulations were performed for 150 ns at constant *T* and *P* (300 K and 1 atm) following standard protocols (46). Only the last 10 ns sampled every 2 ps were used for the subsequent analysis with the idea to evaluate how similar the final state was to the original crystallographic structure, which is the PDB entry 1IHF. It is worth noting that 5J0N was obtained via CryoEM and posterior fitting based on 1IHF. The specific objective of this simulation was to assess the appropriateness of the chosen initial bound state to model the process of DNA wrapping around IHF by reaching a comparable structure to X-ray diffraction.

Free MD simulations of the full DNA construct were done in implicit solvent for a direct comparison with AFM images. The 1λ306 -/+IHF was solvated using the implicit generalized Born model (47) at a salt concentration of 0.2 M with GBneck2 corrections, mbondi3 Born radii set and no cut-off for a better reproduction of molecular surfaces, salt bridges and solvation forces (48, 49). Langevin dynamics was employed for temperature regulation at 300 K with a collision frequency of 0.01 ps^−1^ which reduces the effective solvent viscosity and, thus, accelerates the exploration of conformational space. Prolines were kept intercalated by restraining the distances between key atoms in the proline side chain and the neighboring bases. Following minimization and equilibration, four independent replica simulations of 50 ns were performed of the 1λ306+IHF construct starting from the same minimized structure. One replica of the naked construct was also run for 50 ns starting with a straight B-form DNA. The last 45 ns sampled every 2 ps were used for the subsequent analysis with the objective to characterize the different meta-stable states along the pathways of wrapping the DNA around the protein.

### Analysis of simulations

DNA bend angles were measured for the four independent replicas containing IHF in implicit solvent using the software combo WrLINE/SerraLINE (36, 50). The DNA helix axis was calculated for each frame using WrLINE (50). Then, SerraLINE was used to project WrLINE molecular contours onto the best-fit plane to approximate the experimental methodology (kintetic trapping in 2D) and calculate bend angles (36). The bend angle was defined as the angle between vectors joining points 30 bp apart along the helix axis, where these vectors were separated by a further 30 bp centered on the binding site. SerraLINE and WrLINE are freely available at the repository github.com/agnesnoy. Following the observation of three populations in bend angles, all frames were clustered using hierarchical agglomerative clustering in cpptraj (average-linkage, using the RMSD between frames as the distance metric) until three clusters were formed.

Hydrogen bonds were determined using cpptraj with a distance cutoff of 3.5Å and an angle cutoff of 120°, and the time-average number of intermolecular hydrogen bonds involving each residue was calculated. In all plots, this value was capped at 1, so time-averages could indicate the ratio of frames from the simulation presenting interaction. The hydrogen bonds present in the crystal structure were determined in the same manner based on the minimized 1IHF structure (10), so can only take integer values.

### Umbrella sampling simulations in explicit solvent of DNA wrapping around IHF

We performed a series of umbrella sampling (US) simulations in explicit solvent to accurately calculate the free energy landscape of DNA wrapping around the lateral sides of IHF. The initial structure was the same as used for previous simulations in explicit solvent (*i.e*. unbent DNA molecule of 61 bp bound to IHF just through its protruding β–ribbon arms) and it was prepared as before. The reaction coordinates for the left- and right-hand sides were chosen as the distances between Cα atoms of representative amino acids from the protein interacting far sites (Pro18β and Ser19α, respectively) and the phosphorus atoms from the closest DNA base in the crystal structure. The reaction coordinates were reduced over a series of 5ns simulations in 2Å increments from their positions in the minimized structure (Figure 1A) until the PMF was observed to increase, resulting in a total simulation length of 150 ns for the left arm and 195 ns for the right arm. The final frame of each US window was used as the starting structure for the next. The Grossfield implementation of the weighted histogram analysis method (WHAM) (51) was used to extract the potential of mean force (PMF).

To account for shifts in the free energy landscape between simulations due to the flexibility of the two-atom reaction coordinate, a linear fit was performed in gnuplot to translate the original distances to the distance between the centers of mass of the protein and the 10 bp region of DNA in closest contact with each far side (Figure S4A). The free-energy offsets were estimated by comparison of the means in the plateau of the unbent state.

For each arm, two sets of US simulations were performed. In the first, the other arm was unrestrained and allowed to bind to the protein. In the second, the other arm was held away from the protein by a potential with a spring constant of 2 kcal mol^−1^ Å^−2^ if its reaction coordinate fell below 40 Å. The results from each pair of US sets were considered to represent the two extremes of the other arm’s position, and linear interpolation was performed between them. A two-dimensional landscape was then constructed by simple addition of energies. This was translated into a probability landscape via the function *P*(*F*) = exp(-*F*/*k*_B_*T*). The area representative of each cluster was mapped onto the free-energy landscape by locating each snapshot of the implicitly solvated simulations on the two-dimensional surface using the value of the two reaction coordinates (see Figure S4B). The relative probabilities of the clusters were calculated by weighted integration over the probability landscape when computed with 1 Å resolution, and the probabilities were normalized to sum to 100%.

### Restrained MD simulations in explicit solvent of DNA bridging by IHF

To investigate bridge formation, a second 61 bp piece of DNA was pushed towards the initial IHF-DNA complex formed by unbent DNA of 61 bp bound just to the central part of the protein. This was positioned to lie perpendicular to the main double helix in order to prevent repulsive interactions involving lateral DNA; other arrangements were not as efficient in inducing realistic bridging for this reason (Figure S5). The distance between the backbone atoms closest to their centers of mass (an oxygen atom from a phosphate group and Phe81α Cα) was gradually decreased using a one-sided harmonic potential with a spring constant of 2 kcal mol^−1^ Å^−2^. The asymmetric artificial potential, which was flat for nearer distances but harmonic for farer distances in relation to the target value, was used to avoid biasing of the final bridged structure and, at the same time, to avoid drifting of the second DNA strand away from the protein-DNA complex. Following the formation of a bridge between the two segments of DNA, US as described above was used to pull the second piece of DNA away from the protein, with the reaction coordinate increased until a plateau was observed in the PMF. In all cases, the system was explicitly solvated and set up as before.

## Results

### Modeling the process of DNA wrapping around IHF

We created a structure with the non-curved H2 binding site attached to IHF via only its protruding β-ribbon arms to simulate how DNA becomes wrapped around the protein following the formation of an initial bound state (Figure 1A). We embedded the complex in a DNA construct just over 300 bp for simulations in implicit solvent and subsequent comparison with AFM images over the analogous construct. A shorter 61 bp DNA fragment was extracted from the long piece of DNA with the idea of performing a more rigorous simulation in explicit solvent that would evaluate the validity of the implicit solvent model and the chosen initial bound state (see Materials and Methods). A continuum representation of the solvent reduces the computational cost of simulations compared with the use of an actual solvation box with discrete water molecules and ions, allowing the size of the system to be scaled up to what is workable for AFM experiments. However, non-bonded interactions are not so accurately described on implicit solvent, especially those based on electrostatics (52), so caution is always needed (53).

We started free MD simulations from the open state shown in Figure 1A using both solvent models, and allowed the DNA to spontaneously wrap around IHF, as in Figure 1B. Time-evolution of the root mean square deviation with regards to the X-ray diffraction structure 1IHF (Figure 1C) shows that wrapping close to that observed in the crystal structure is obtained for both types of solvent without a significant difference in their converged stage (RMSd is 4.79 ± 0.54 Å and 4.98 ± 0.16 Å in explicit and implicit solvent, respectively). The observed DNA–protein interactions correspond well to those present in the 1IHF crystal structure (10) (Figure 1D), including the insertion of Ser47α and Arg46β residues into the minor grooves of lateral DNA (Figure 1B) and the additional interactions between the DNA backbone and positively-charged and polar amino acids.

These interactions can be broadly divided into four regions based on their position relative to the center of the binding site and the protein subunit to which the involved amino acid belongs. On the left-hand side (containing the A-tract), the α subunit is closer to the center and thus constitutes the “near left” side, while the β subunit is farther and composes the “far left”. On the right-hand side (containing the consensus sequence), the α and β subunits are inversely arranged, delimiting the “far right” and “near right” sides, respectively (see Figure 1B).

There is generally strong agreement between the DNA-protein hydrogen bonds presented by the X-ray diffraction structure and simulations irrespective of the solvent model used. We observed a more defined set of interactions at the “near” sites compared with the “far” ones that could be caused by an increase of flexibility on the DNA part farther away from the main recognition side. In addition, the shortness of the DNA at the crystal structure (35 bp versus 61 bp on simulations) makes it difficult to capture some of the more distant interactions.

In general, our results demonstrate the validity of the molecular dynamics methodology for this system, as well as suggesting an explanation for the previously proposed two-step binding mechanism. In this two-step model, the IHF arms would bind to DNA first and then the proline residues would intercalate to induce flexible hinges prior to wrapping. Although our simulations start from straight DNA with the prolines already intercalated, it is possible that some bending may occur following the initial recognition that would facilitate the intercalation of prolines and the subsequent strong kink. In any case, the binding mechanism that we propose explains both the high activation energy of initial binding (27) due to proline intercalation and the smaller free energy of wrapping (28).

### Multimodality of IHF specific and non-specific binding revealed by AFM

To investigate the differences between specific and non-specific binding, two short sequences were amplified from phage λ, one with no IHF consensus sequence (0λ361) and another with a single consensus sequence (1λ306) of lengths 361 bp and 306 bp respectively (Figure 2). The lack of sequences with similarity to the IHF specific recognition motif was confirmed by scoring the entire 0λ361 molecule to the PSSM available at the Regulon database (33) (see Figure 2, S1 and Materials and Methods). The two DNA fragments were then imaged using AFM and compared to MD simulations to determine the different morphological behaviors.

DNA contours were recovered by skeletonizing pre-processed AFM images (see Materials and Methods), with qualitatively different behavior observed depending upon the presence of IHF (Figures 3). Figure 3E shows a limited reduction of the radius of gyration for the two constructs if IHF is present, suggesting that the protein bends DNA to some extent in both cases, in a specific manner in the case of 1λ306 and non-specifically for 0λ361. However, as the radius of gyration and the rest of dimensional parameters (Figure S2) can only determine overall compaction, a more precise bending angle analysis was also performed.

**Figure 3.**
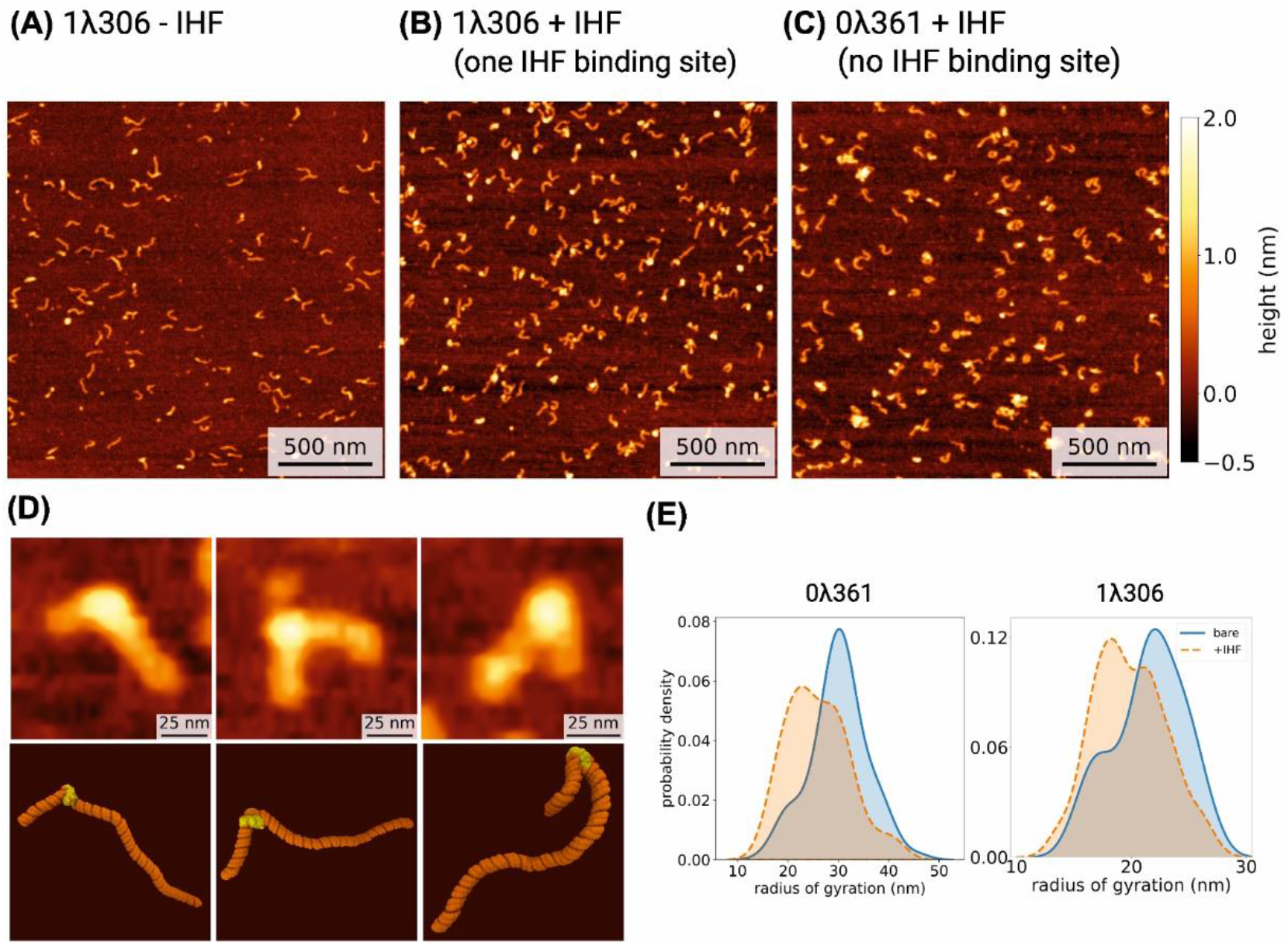
Atomic force microscopy (AFM) images of DNA with and without IHF. **(A)** AFM image of DNA with one IHF binding site (1λ306) without IHF. In the presence of IHF (1IHF:5 bp), DNA with one IHF binding site (1λ306) **(B)** shows more binding when compared to DNA without a binding site (0λ361) **(C)**, although globular artefacts from poly-L-lysine can be seen. **(D)** Three representative examples of IHF binding, with a comparison of AFM (top) and molecular dynamics simulations (bottom). **(E)** Kernel density estimates of the radius of gyration of DNA ± IHF.

Our limited resolution prevented us from measuring DNA angles around IHF exclusively, so we considered bending distributions of naked DNA as a background (see Materials and Methods). A large proportion of the total values (^~^60-70%) obtained with IHF were indistinguishable from the bending behavior of bare DNA molecules (Figures S3A-B) as a result of the natural overlap between the two angle distributions and the presence of some unbound DNA, and this was accounted for in the subsequent analysis.

The addition of IHF leads to a further two and three peaks in the angular distributions for 0λ361 (Figure 4A) and 1λ306 (Figure 4B), respectively, compared to the single peak found in bare DNA, according to the reduced chi-square goodness-of-fit tests (p-values >0.99; see Figures S3C-D). The two-sided Kolmogorov-Smirnov test was applied to confirm the statistical difference of 1λ306 bend distribution compared to 0λ361 in the presence of IHF (p-value=0.002) and from naked DNA (p-value=0.008). The same test did not find statistically significant difference between distributions of 0λ361 with/without IHF, probably due to the larger overlap and the larger amount of unbound DNA caused by the relatively low affinity, although the presence of more than one peak was suggested by the previous chi-squared test.

**Figure 4.**
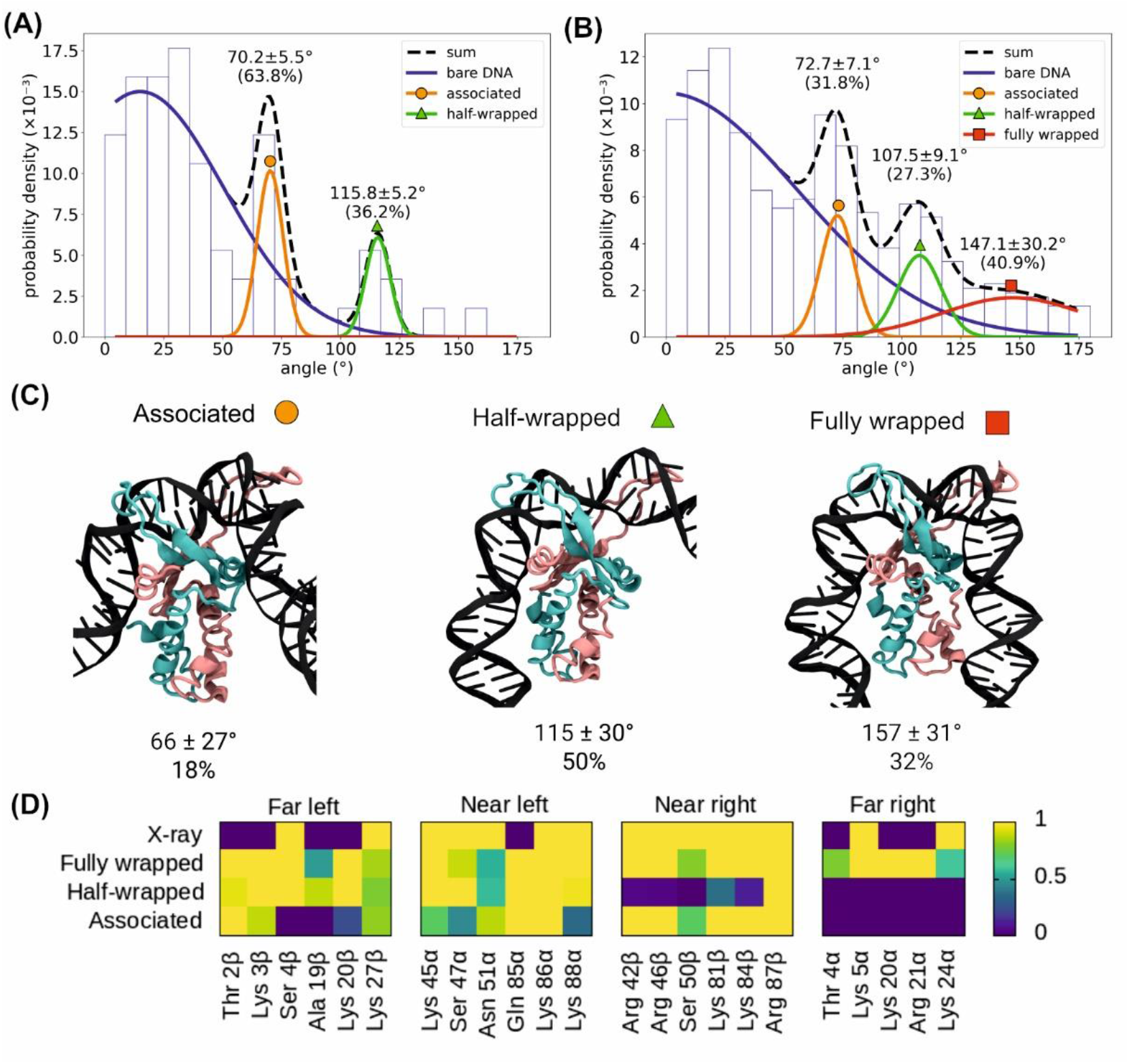
Analysis of different bending angles of IHF-bound DNA. Bending angle distribution obtained by AFM of 0λ361 (n=87) **(A)** and 1λ306 (n=144) **(B)** with IHF. Distributions observed without IHF (blue lines, Figures S3A-B) are considered as background and disregarded. The remaining peaks are well fitted by Gaussians (as shown by the reduced *χ^2^* in Figures S3C-D), representing two common states with bending angles around ^~^70° and ^~^110° and an additional state presented only by 1λ306 with an angle around ^~^147°, closer to the X-ray structure. The broadness of the peak around ^~^147° is caused by resolution issues that make angles more difficult to measure as they get larger (23). **(C)** Simulations of the same construct (in implicit solvent) can be classified into three clusters, the mean bending angles of which correspond well with the experimental data. Proportions for each state are obtained via US simulations in explicit solvent (see Figure 5). Colour scheme is the same as in Figure 1. **(D)** The clusters (from implicitly solvated simulations) can be characterized by the DNA–protein hydrogen bonds present in each state; while the fully wrapped DNA interacts with the protein on both sides, the half-wrapped DNA interacts only on the left, and the associated state interacts primarily with the “near” sites. Contact maps show the average number of frames presenting at least one hydrogen bond between DNA and each residue using all frames belonging to each cluster or from the 1IHF crystal structure.

The peaks around 73 ± 7° and 107 ± 9° (mean ± s.d. values from 1λ306) are common to both constructs, suggesting that they occur due to non-specific binding, whilst the canonical large bending angle (147 ± 30°), as seen in the crystal structure (5), only appears in 1λ306 that contains one specific IHF binding site. The proportions of the 73°, 107° and 147° states (approximately 32%, 27% and 41% respectively) show that all three binding modes are present in roughly equal quantities for the construct with a specific binding site (1λ306). However, the proportion of the 73° state (^~^64%) to the 107° (^~^36%) state is much larger for 0λ361, suggesting the former state is mostly related to non-specific binding. Some binding could still occur as similar sequences to the left part of the specific binding site (an A-tract followed by AT base pairs downstream) are present in the sequence of 0λ361, although they do not count towards the PSSM calculation (Figure S1).

We then isolated the subpopulation of DNA molecules with bend angles >54° and >126° in the presence of the protein and we observed a reduction (mean ± s.e.) in the radius of gyration (19.1 ± 0.2 nm and 17.3 ± 0.4 nm, respectively) and end-to-end distance (51.5 ± 0.8 nm and 37.0 ± 1.5 nm, respectively) compared with the construct in the absence of IHF (radius of gyration of 22.4 ± 0.4 nm and end-to-end distance of 68.5 ± 1.4 nm), demonstrating the global effect of the IHF-induced bend. Our results are largely consistent with previous AFM studies where a broad distribution of angles was detected between approximately 80-150° (18, 19). The 73° state is one that has not been observed in AFM before, possibly as it is within the expected range of angles for bare DNA and could be excluded if not accounting for the bare DNA distribution (Figures S3A-B) or due to the selection of an H2 binding site.

### The multiple IHF binding modes are confirmed by MD simulations

To validate the states deduced from AFM, four independent (replica) unrestrained MD simulations were performed of the 1λ306+IHF construct in implicit solvent (Figure 3D, Supplementary Movies 1-4), starting from the bound but unbent state shown in Figure 1A. One unrestrained MD simulation was also performed of the naked 1λ306 molecule in implicit solvent for comparison. We imitated the kinetic trapping of DNA molecules to the surface observed in our experiments by projecting the simulations to the best fitted plane so we could compare DNA bend angles (see Materials and Methods). Averages and standard deviations of radius of gyration (25±1 nm and 21±1 nm, -/+IHF respectively) and end-to-end distances (81±7 nm and 49±8 nm, -/+IHF respectively) of the projected trajectories were found to be within the experimental error (Figure S2), thus demonstrating the validity of our approach. In addition, we found that the deviation from planarity of the complexes was less than 10% on average, which indicated small distortions on the complex resulting from being trapped in 2D (Figure S6).

All the frames of the replicas with IHF were merged together and were classified into three groups through hierarchical agglomerative clustering using RMSD as the distance metric, finding groups with bend angles of 66 ± 27°, 115 ± 30°, and 157 ± 31°, respectively, in good agreement with the AFM results (see Figure 4). A representative frame was selected for each group in Figure 4C and the mean bend angle was calculated for each. The clusters are also characterized by the hydrogen bonds between the protein and DNA (Figure 4D). In the “fully wrapped” state (with a bend angle of 157°), both sides of the DNA form hydrogen bonds with both subunits of the protein; in the “half-wrapped” state (115°), the left-hand side, which contains the A-tract, forms contacts with both subunits while the right-hand side forms no hydrogen bonds; and in the “associated” state (66°), only the “near” subunits on the left and right sides form hydrogen bonds with the DNA.

Bend angle distribution extracted from the replicas with IHF exhibits three peaks (68±37°, 117±4° and 149±23°; see Figure S4C) that align well with AFM and with the previous structural classification, demonstrating the consistency of our methodology. However, we do not expect good agreement on the proportions of the different states (4% associated, 42% half-wrapped, 54% fully wrapped) due to the limitations on the sampling.

Each individual replica presents a slightly different behavior in terms of the time spent at the different states and in the transitions between them (Figure S4D). In the first one, the complex rapidly reaches the fully wrapped state after briefly passing through the half-wrapped (Supplementary Movie 1). Because this is the replica with the most canonical behavior (more like the crystal structure), it is the one selected in Figure 1 for assessing the recovery of crystallographic interactions by means of implicitly solvated simulations. In the second replica, the complex oscillates between the half- and fully wrapped states (Supplementary Movie 2), and, in the third replica, it remains in the former (Supplementary Movie 3). In the last replica, the complex traverses the associated state before arriving to the fully wrapped state (Supplementary Movie 4). These simulations suggest a metastable nature for the partially bent states (associated and half-wrapped) in the course of DNA binding around IHF before reaching the global minimum occupied by the fully-wrapped state.

To properly describe the free energy landscape of the wrapping process and the proportion of the different states, we performed restrained MD simulations (54, 55) of a 61 bp segment of the construct in explicit solvent (see Materials and Methods). A more accurate representation of the solvent was chosen on this occasion for describing DNA-IHF interactions at a more rigorous level, enabling us to verify the previous implicitly solvated simulations. The distance between the bottom part of the protein and the interacting DNA on each side was varied in a series of US simulations, once with the other arm allowed to wrap and once with the other arm held away from the protein. The PMF for each simulation was calculated using the WHAM method (51, 56). By linearly interpolating between the two sets of results for each arm (Figure 5A-B) a two-dimensional free energy landscape was constructed with the reaction coordinates as orthogonal axes, as in Figure 5C.

The relative probabilities of the clusters were determined by integrating over the relevant regions of the free energy landscape (Figure 5D and Figure S4B). This analysis predicts that the canonical fully wrapped (157°) state should be observed around 32% of the time, the half-wrapped (115°) state 50%, and the associated (66°) state 18%; while these figures do not directly agree with AFM, probably due to the differences between the simulation and experimental conditions, they do predict that all three states should occur in significant proportions. Each cluster corresponds to a minimum or plateau in the free energy landscape; the locations of these features, along with the observed paths between them (Figure S4D), are highlighted in Figure 5D.

### Asymmetry between DNA arms causes a mechanical switch towards strong bent DNA

As discussed in the previous section, it is notable that the consensus sequence does not interact with IHF in the half-wrapped state and this state is also observed in the absence of a specific binding site. The free energy surface, which is described as a function of the two distances between the bottom of the protein and the two DNA arms, reveals that there is a large activation barrier preventing the right-hand side (which contains the consensus sequence) from binding before the left-hand side (which contains the AT-tract) (Figure 5C). The form of the potentials is also qualitatively different. While the left-hand side presents a relatively deep potential with a minimum around 34 Å and an additional small plateau at around 40 Å, implying that binding is energetically favorable (Figure 5A), the right-hand side has a much flatter shape with small minima around 31 Å and 43 Å, and appears to be dominated by thermal noise (Figure 5B).

While the free-energy landscape presented by the left-hand side appears to be independent of the position of the right arm, the right-hand side is prevented from binding fully by a large barrier when the left arm is held away from the protein; this barrier is not present when the left arm is fully bound. This barrier is associated with a change in the structure of the protein. When the left arm is unbound, the upper subunit of the protein on the right-hand side protrudes, so full binding of the right arm would require extra DNA bending apart from the flexible region. Binding of the left arm flattens the surface presented to the right arm, removing this physical barrier and allowing the DNA on the right-arm to bind while remaining mostly straight (Figure 5D).

This free energy landscape suggests that it is very energetically favorable for the protein to bind the A-tract on its left-hand side, but that it is less inclined to bind the consensus sequence, and that the A-tract must bind first (Supplementary Movie S5). This is reflected in the observed clusters—there is no cluster in which the left-hand side is unbound, while the position of the right-hand side distinguishes the fully and half-wrapped states (Figure 4C–D). The associated state corresponds to the smaller local plateau or minimum in each arm’s energy landscape (Figure 5), as seen when the values of the reaction coordinates are plotted together for each frame of a cluster as in Figure S4B.

The lack of interaction of IHF with the consensus bases suggests that the half-wrapped and associated states are forms of non-specific binding, a conclusion supported by the occurrence of bend angles corresponding to these states in AFM observations of the 0λ361 construct (in which the consensus sequence does not appear). The consensus sequence is therefore necessary only for the fully wrapped binding mode, suggesting that initial binding of IHF to DNA occurs without sequence specificity and a consensus sequence enhances bending.

The free energy landscape features a significant minimum that appears to correspond to another state (represented by a white cross in Figure 5D), in which the left-hand side is fully bound and the right-hand side is partially bound. However, the frames in this region are difficult to distinguish from those in the fully wrapped state as the bending angles are very similar, and the corresponding minima in the free-energy landscape are separated by only a narrow energy barrier of less than 1 kcal/mol. Since the system can move freely between these two minima under the influence of thermal noise, it is more appropriate to consider this state an extension of the fully wrapped mode.

The asymmetric cooperative behavior observed between arms in the presence of a specific sequence could serve for reinforcing the prevalence of the fully-wrapped state, as it reduces the amount of accessible intermediate states that the system would present in the case of both arms moving in a totally uncorrelated manner. This interconnection between the two halves would help the IHF to sharply bend the DNA with a substantial probability in a similar way to a mechanical switch.

### DNA aggregation and bridging by IHF

At concentrations of 1 IHF:5 bp, occasional small clusters of DNA/protein could be seen for both 0λ361 and 1λ306 (Figures 6). Further increases of the IHF concentration causes the formation of clusters for 1λ306 (Figure 6C). This behavior at a higher concentration (^~^1 IHF:3 bp) has previously been seen (19) and shows how large cellular concentrations of IHF – such as in the stationary phase – could also increase genome compaction by bridging.

**Figure 5.**
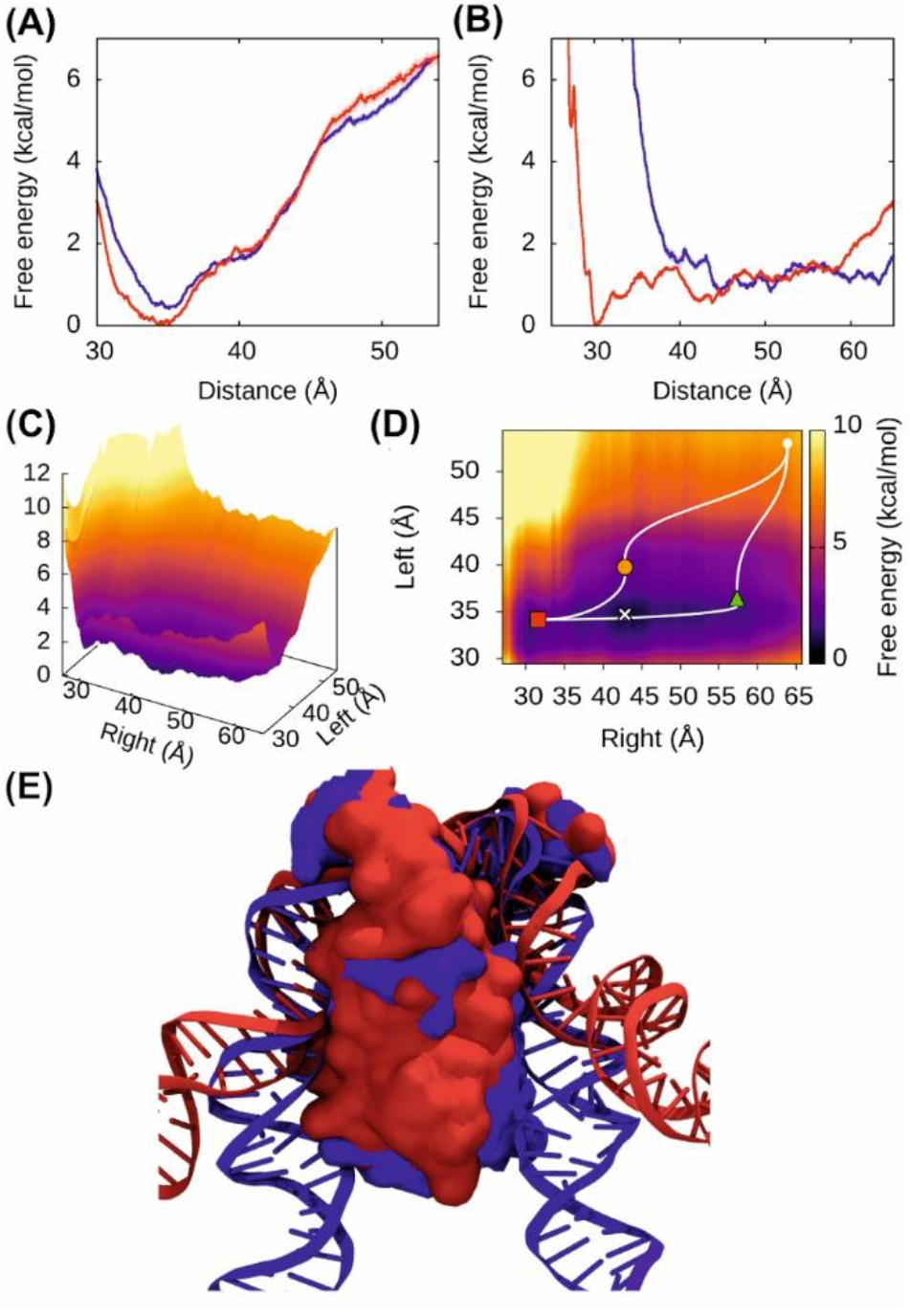
Free energy of DNA bending by IHF. The free energy landscapes determined through US simulations in explicit solvent differ for the left arm **(A)**, containing the A-tract, and the right arm **(B)**, containing the consensus sequence. The left arm presents a deep potential well regardless of whether the right arm is free to bind (red) or held away from the protein (blue), while the right arm can bind fully only when the left arm is bind (red) and not held away (blue). **(C)** The 2D reconstruction of the energy landscape via linear interpolation between arm’s positions shows the striking asymmetry. **(D)** The observed clusters from implicitly solvated simulations correspond to minima and plateaus in the free energy landscape plotted here along with the approximate paths by which replicas traversed. Fully wrapped is represented by a red square, partially-wrapped by a green triangle and associated state by orange circle. The white cross represents a sub-state from the fully wrapped form where left-hand side is fully bound and the right-hand side is partially bound. **(E)** Two superimposed structures with the left arm bound (blue) and unbound (red) showing that, when the left arm is unbound, the upper subunit protrudes on the right-hand side, making it difficult for the DNA to interact with the bottom of the protein.

**Figure 6.**
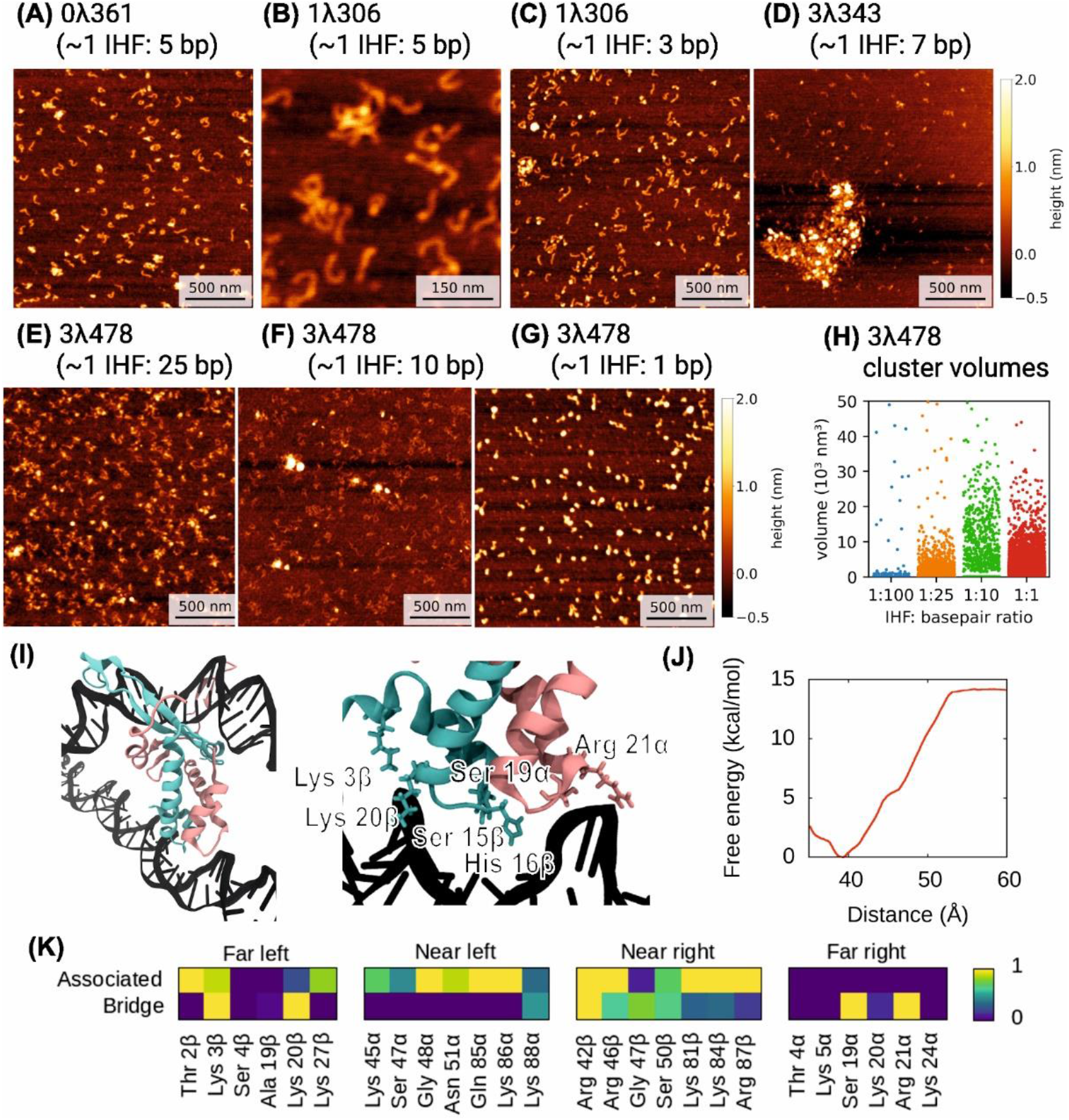
IHF-bound DNA forms aggregates depending on protein concentration and on the type of DNA constructs. A large amount of IHF leads to some aggregation on 0λ361 **(A)** and 1λ306 **(B-C)**, with no or one binding site respectively. DNA constructs with 3 binding sites (3λ343 and 3λ478) start to form clusters at lower concentrations. **(D)** Large aggregates are observed for 3λ343. **(E-F)** 3λ478 forms more numerous and/or denser clusters as IHF concentration increases until saturation is achieved **(G). (H)** Cluster volumes for 3λ478 as function of IHF concentration **(I)** Bridge structure and zoom from MD simulations with the interacting amino acids highlighted in atomic representation. Colour scheme is the same as in Figure 1. **(J)** Free-energy landscape of bridging formation relative to an unwrapped state with a close additional DNA strand (Figure S5A). **(K)** Contact maps are calculated as in Figures 1 and 4 revealing that the residues from the “far” interacting regions are those which form the bridge. The DNA strand into which IHF intercalates remains mostly unbent, interacting with the protein even less than in the associated state.

A construct with three binding sites (3λ478) behaves very differently to the previous constructs that contain one or no specific binding sites. Even at low concentrations, from around 1 IHF: 10 – 25 bp (Figures 6E and 6F), a greater number of IHF/DNA clusters appear to form with a volume of the order of 10^4^ nm^3^ (Figure 6H) compared to volumes of around 10^3^ seen for individual double-stranded DNA molecules (Figure S2). At larger concentrations (1 IHF: 1 bp), IHF was numerous enough to coat the DNA reducing the size of visible clusters (Figure 6G-H).

A shorter form of this construct (3λ343, 343 bp long) (Figure 6D) was seen to show larger aggregates, reaching volumes of order 10^5^-10^6^ nm^3^ at 1 IHF:7 bp. The reason for the prominence of aggregation on the latter construct could be related with the elimination of spare DNA, which might reduce electrostatic repulsion and steric extrusion between DNA duplexes. 3λ343 and 3λ478 are highly homologous apart from a tail of around 125 bp on one side of the three binding sites in the longer construct that has a much lower AT-content (52%) compared with 3λ343 (70%) and thus presents lower affinity to IHF (Figure S1). Although these conglomerates may be due to protein aggregation, the difference between these and the earlier constructs demonstrates that it is not the case.

IHF-induced aggregation can be explained by the bridging of two separate double-stranded DNA segments by a single IHF dimer, a phenomenon observed in our MD simulations (Figure 6I). These bridges result from non-specific interactions involving positively charged and polar amino acids from the far binding sites of the protein and the backbone of DNA (Figure 6K). US simulations (see Materials and Methods) reveal that bridging is very energetically favorable when a second double-stranded DNA is close enough, with a free energy difference on the order of 14 kcal/mol between the bridged and unbridged states (Figure 6J). The main DNA fragment, into which the prolines intercalate, remains mostly unwrapped, probably due to the electrostatic repulsion exerted between the two double-stranded DNA molecules (Figure 6I and S5).

The combination of AFM and MD suggests that the clustering of constructs containing three binding sites is due to the capacity of IHF to bind more than one DNA strand at the same time. We observe that DNA·IHF·DNA bridging is favourable even for low concentrations of IHF. The strong electrostatic repulsion between close DNA strands could prevent the complete wrapping of DNA around the protein, even on sequence-specific sites, facilitating the bridging interaction described above.

## Discussion

Here we have integrated advanced theory and experiments from physics to explore a basic phenomenon of life at the single-molecule level (57), namely how individual proteins interact with DNA. Our findings enable new insights into the complex interactions between proteins and DNA. Both AFM experiments and MD simulations show evidence of a larger number of bending modes of the IHF/DNA complex than previously proposed. The presence of three states (fully wrapped, half-wrapped and associated) as well as the large proportion of unbound DNA suggests a multi-modular system, where higher AT-content makes binding more probable (58, 59) but in which a consensus sequence is needed to fully bend DNA. We put forward a model for the mechanism of IHF binding where full wrapping is sequence specific but initial binding is not (Figure 7).

**Figure 7.**
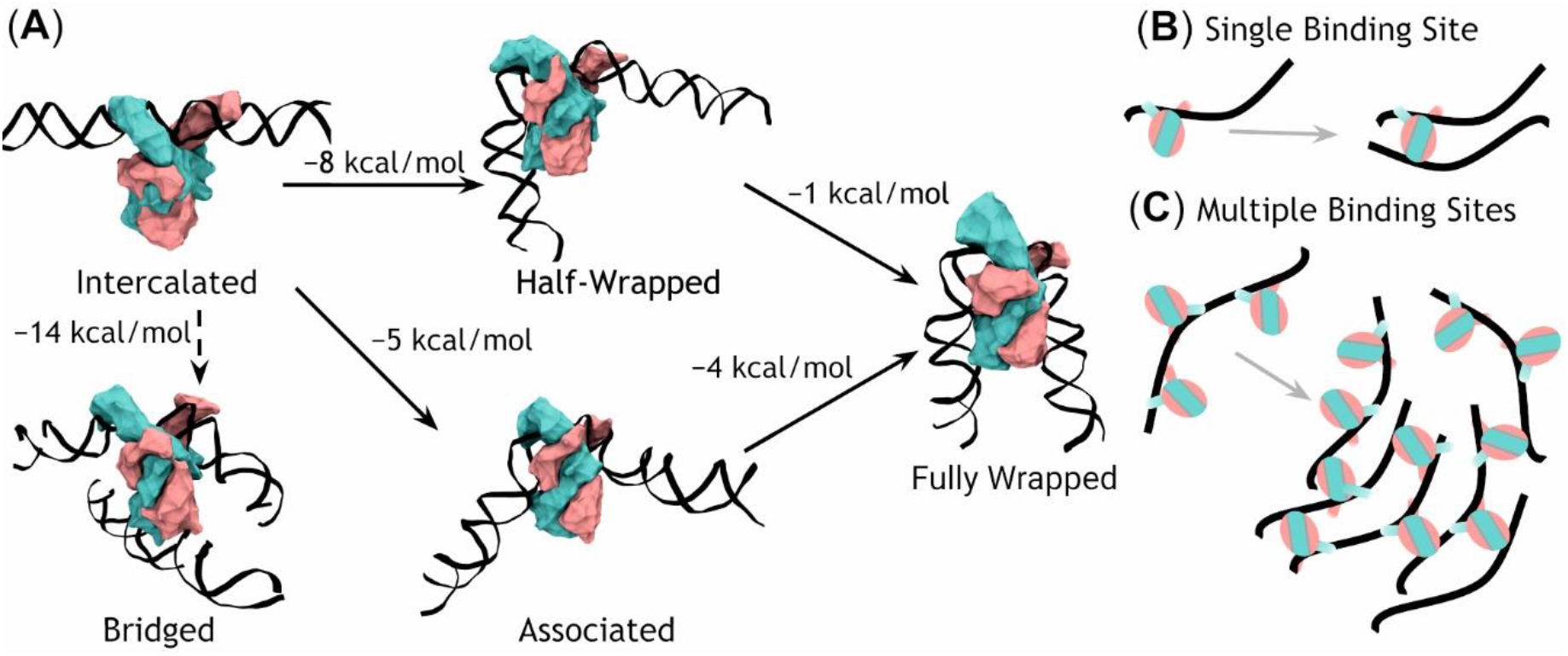
Model of IHF binding – bending and bridging. **(A)** IHF first recognizes DNA through its β-arms and the prolines intercalate into a double-helix structure that its straight in our model although it might already contain a certain bent. The first binding step appears to be either a loose association to the DNA (73° binding) or a half-wrapped state (120°). Both can then progress to the canonical fully wrapped state. As the initial state leaves the bottom half of IHF unbound it can also bind to another strand of DNA non-specifically resulting in a bridge (transition conditioned to the second DNA strand being nearby, dashed arrow). Free energy differences between states (ΔF) were estimated via US simulations in explicit solvent. **(B)** A construct with a single binding site can only form a single bridge whereas **(C)** multiple binding sites can form multiple bridges, leading to aggregation.

First, we provide structural insights into the two-step binding mechanism previously proposed (27), as the crystallographic structure is reached by our simulations when started from a bound but unwrapped DNA (Figure 1). Thus, our results reinforce the idea that the protein first binds DNA non-specifically via its extended arms; the prolines at the tips of the arms would then intercalate with an activation energy around 14 kcal/mol (27); from there, DNA wraps around the protein in a relatively downhill process stabilized by up to 9 kcal/mol, a value calculated from our simulations. This is a larger estimation compared to single-molecule experiments (3.6 kcal/mol) (28) because simulations were started with the most possible unbent configuration for DNA, so it is the upper limit of free energy relaxation associated to the wrapping process. Either the intercalation of prolines is followed by a rapid bending relaxation, or the first binding step favors some bending that facilitates proline intercalations, being the two contributions (intercalation or bending) difficult to be neatly discriminated from experimental data. In any case, simulations support a bind-then-bend mechanism that has also been observed in other DNA-flexing proteins (60).

When IHF binds to DNA, it is highly likely that the left-hand side will bind to the DNA first, but not with the consensus part of the sequence. This results in the half-wrapped state, which is ^~^8 kcal/mol more stable than the initial structure and in which the left-hand side is fully bound while the right-hand side is free. Alternatively, both sides may make interactions with the nearest subunit of the protein, resulting in a smaller bending angle, that we designate the associated state (favored by ^~^5 kcal/mol). Transitions between the associated and half-wrapped states were not observed (Figure S4D), and both appear to be long-lived metastable states corresponding to plateaus or local minima in the free-energy landscape (Figure 5D). As the associated and half-wrapped states are experimentally seen in constructs both with and without a consensus sequence, this suggests a non-specific binding mechanism for these states. However, in the presence of a consensus sequence, both of intermediate states lead towards the global minimum (Δ*F* ≈ −9 kcal/mol relative to the initial structure), resulting in canonical binding.

Previous experiments (19, 29, 30) already indicated that IHF could bind DNA in more than one state, generally in two states which were broadly described as a fully and partially bent. Magnetic tweezers experiments appeared to show a smaller angle state ^~^50° (19) (i.e. similar to our associated state). The lack of detection of a third state could be due to the fact that the force applied on the DNA was greater than the few tenths of piconewtons we predict is needed to overcome the potential barrier between the half-wrapped and fully wrapped states. By using fluorescence-lifetime-based FRET (29), Ansari and co-workers deduced the presence of three binding modes and that two of these involved partially-bend DNA, in line with the results presented here, although they did not have the resolution and/or the associated modelling to detect the structural properties of the non-canonical conformation. The proportion of the fully-wrapped state was found to be higher in their experiments than in our results, but this difference might be caused by the use of a different binding site (H’ vs H2, see Figure 2).

The half-wrapped and associated states may not be the only conformations in which IHF binds non-specifically to DNA. Hammel *et al*. (61) obtained crystal structures of HU where the DNA was bound across the α-helical body of the protein rather than between the extended β-ribbon arms. The similar electrostatic profiles on IHF and HU (see Figure S7) suggest that this extra non-specific binding mode could be possible in IHF. However, such state would need a different initial conformation for being explored by simulations and we predict that it would not induce a significant bend stronger than the typical angles for bare DNA, so this possibility was not explored in this work.

The asymmetric allostery, by which DNA binds to the right side only after binding to the left side, makes the fully wrapped state relatively more probable than both arms moving independently. This mechanism, thereby, constructs a mechanical “switch”, since the ultimate structural configuration can switch between different states in a mechanically dependent manner. The formation of such a sharp bend in one of these observed states is impossible to achieve in naked DNA, even for the most curved sequences (62, 63) unless base stacking and complementary hydrogen-bond pairing are disrupted on the double helix generating kinks or melting bubbles (36).

This regulatory behavior could be switched on or off by tension, structural influences upstream of the binding site or by different levels of DNA supercoiling. The semi-stable states observed would allow IHF to remain associated with the DNA while retaining some flexibility, which could be important in the formation of higher-order nucleoprotein complexes such as transcription regulatory loops. Similarly, other proteins in mammalian systems might expect to behave in similar meta-stable ways when interacting with DNA under physiological levels of superhelical tension. For example, nucleosomes present spontaneous unspooling of the outer stretches of DNA causing multiple levels of wrapping around the histones (64). These meta-stable states are modulated by tension (65) or by the presence of neighboring nucleosomes (66) and regulate the access to nucleosomal DNA (67).

Remarkably, we observe that the more conserved bases are within the region that is more dynamically bound to IHF, which could illustrate the need for flexibility downstream of the protein or, as an alternative, it could reflect the difficulty of identifying consensus sequence motifs, and therefore cognate binding sites, in the case of DNA shape recognition (68). Similarly, it was found that the transcription factor GabR from *Bacillus subtilis* needs to recognize flexed DNA at a location distinct from its known binding sites (69).

Another interesting behavior observed was the bridging/clustering of DNA by IHF. At high concentrations of IHF (such as during periods of inactivity of cell division and DNA replication when the bacterial nucleoid is at its most compact in the cell cycle) this can occur non-specifically, giving it a role in compaction. On the other hand, the constructs with multiple binding sites seem to preferably select for the bridging behavior over bending when DNA molecules are moderately concentrated. This behavior shows how the formation of bridges could be driven by the screening of electrostatic repulsion between neighboring DNA molecules thanks to positively-charged architectural proteins (70). In this regard, hidden secondary recognition sites causing DNA bridging by means of basic amino acids have also been identified for other bacterial architectural proteins like Topoisomerase IB (71) and ParB (72, 73). This phenomenon could also be promoted by steric hindrance or tension due to the presence of other proteins preventing complete wrapping by each IHF, leaving the lower regions of the protein free to bind other DNA

Our study not only explains the role of IHF in biofilms, by cross-bridging extracellular DNA, but undoubtedly shows that the function of IHF is far more multifaceted than bending DNA. We observe that specific binding sites can be simply modulated or extended by additional non-specific electrostatic-driven interactions between the protein and the DNA (70, 74). In fact, positively charged patches on DNA-interacting proteins have been traditionally used for predicting DNA-binding interfaces (75). We anticipate that this could be a general mechanism used by other NAPs (76) and eukaryotic chromatin-binding proteins (77, 78) to enable a variety of DNA bending and bridging modes. Promiscuous electrostatic interactions between negatively-charged DNA and positively-charged genomic architectural proteins could be one of the primary molecular forces underpinning the physical organization of all kinds of chromosomes, including the formation of membrane-less phase-separated condensates inside cells (79–81). Finally, the general methods we have developed here for comparing AFM imaging with MD simulations have a utility that could be applied for many other protein-DNA interactions in the chromosome, beyond just IHF.

## Data Availability

All relevant data is included in the main manuscript, the supplementary material and the University of York Data Repository (https://doi.org/10.15124/0b602b82-16e9-4fa8-9a64-a15d7373f80e)

## Acknowledgements

We thank the Physics of Life (PoL) Group, University of York for providing pump-priming resources. This work was supported by the Engineering and Physical Sciences Research Council (EPSRC) [EP/N027639/1, EP/T002166/1, EP/R029407/1 and EP/P020259/1]; Biology and Biotechnology Research Council (BBSRC) [BB/R001235/1]; Leverhulme Trust [RPG-2017-340]. V. V.-B. was funded by CONACYT agency from Mexican government (scholarship no 291163). Calculations were performed on ARCHER, JADE, Cambridge Tier-2 and the local York facilities (Viking and YARCC clusters). We would also like to thank Alice Pyne for her assistance with the AFM experimental techniques and analysis.

