## supplementary methods for "Integration host factor bends and bridges DNA in a multiplicity of binding modes with varying specificity"

### Supplementary Information

#### Supplementary Tables and Figures

**Table S1. Primers for DNA amplification by PCR of sequences used, amplified from phage λ.**

| Primer Name | Sequence |
|---|---|
| 0λ361-F | 5`- GCATCATCAAGTGCCGGTCG -3` |
| 0λ361-R | 5`- TGCTGTTGGTTGCACTGCTG -3` |
| 1λ306-F | 5`- CAAGACACCGGATCTGCAC -3` |
| 1λ306-R | 5`- GCATATGATGTCTGACGCTGG -3` |
| 3λ343-F | 5` - CTTTGTGCTTCTCTGGAGTGCG – 3` |
| 3λ343-R | 5` - GGCAGGGAGTGGGACAAAATTG – 3` |
| 3λ478-F | 5` - GATTGCGAGGCTTTGTGCTT - 3` |
| 3λ478-R | 5` - CTACCTTTCACGAGTTGCGC - 3` |

**(A)**

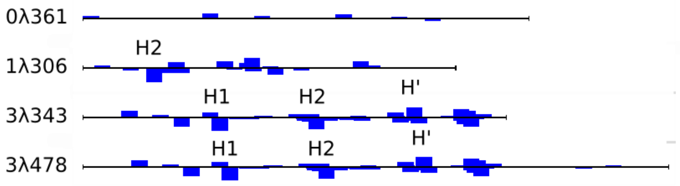


**(B)**

0λ361
GCATCATCAAGTGCCGGTCGTGCAGCTTCCTCGGCAACGGCGGCAGAAAA -50
TTCTGCCAGGGCGGCAAAAACGTCCGAGACGAATGCCAGGTCATCTGAAA -100
CAGCAGCGGAACGGAGCGCCTCTGCCGCGGCAGACGCAAAAACAGCGGCG -150
GCGGGGAGTGCGTCAACGGCATCCACGAAGGCGACAGAGGCTGCGGGAAG -200
TGCGGTATCAGCATCGCAGAGCAAAAGTGCGGCAGAAGCGGCGGCAATAC -250
GTGCAAAAAATTCGGCAAAACGTGCAGAAGATATAGCTTCAGCTGTCGCG -300
CTTGAGGATGCGGACACAACGAGAAAGGGGATAGTGCAGCTCAGCAGTGC -350
AACCAACAGCA

1λ306
CAAGACACCGGATCTGCACAACATTGATAACGCCCAATCTTTTTGCTCAG -50
ACTC**TAA**CTCA**TTGATA**CTCATTTATAAACTCCTTGCAATGTATGTCGTT -100
TCAGCTAAACGGTATCAGCAATGTTTATGTAAAGAAACAGTAAGATAATA -150
CTCAACCCGATGTTTGAGTACGGTCATCATCTGACACTACAGACTCTGGC -200
ATCGCTGTGAAGACGACGCGAAATTCAGCATTTTCACAAGCGTTATCTTT -250
TACAAAACCGATCTCACTCTCCTTTGATGCGAATGCCAGCGTCAGACATC -300
ATATGC

3λ343
CTTTGTGCTTCTCTGGAGTGCGACAGGTTTGATGACAAAAAATTAGCGCA -50
AGAAGACAAAAATCACCTTGCGCTAATGCTCTGTTACAGGTCACTAATAC -100
CATC**TAA**GTAG**TTGATT**CATAGTGACTGCATATGTTGTGTTTTACAGTAT -150
TATGTAGTCTGTTTTTTATGCAAAATCTAATT**TAA**TATA**TTGATA**TTTAT -200
ATCATTTTACGTTTCTCGTTCAGCTTTTTTATACTAAGTTGGCATTATAA -250
AAAAGCATTGCT**TATCAA**TTTG**TTG**CAACGAACAGGTCACTATCAGTCAA -300
AATAAAATCATTATTTGATTTCAATTTTGTCCCACTCCCTGCC

3λ478
GATTGCGAGGCTTTGTGCTTCTCTGGAGTGCGACAGGTTTGATGACAAAA -50
AATTAGCGCAAGAAGACAAAAATCACCTTGCGCTAATGCTCTGTTACAGG -100
TCACTAATACCATC**TAA**GTAG**TTGATT**CATAGTGACTGCATATGTTGTGT -150
TTTACAGTATTATGTAGTCTGTTTTTTATGCAAAATCTAATT**TAA**TATA**T** -200
**TGATA**TTTATATCATTTTACGTTTCTCGTTCAGCTTTTTTATACTAAGTT -250
GGCATTATAAAAAAGCATTGCT**TATCAA**TTTG**TTG**CAACGAACAGGTCAC -300
TATCAGTCAAAATAAAATCATTATTTGATTTCAATTTTGTCCCACTCCCT -350
GCCTCTGTCATCACGATACTGTGATGCCATGGTGTCCGACTTATGCCCGA -400
GAAGATGTTGAGCAAACTTATCGCTTATCTGCTTCTCATAGAGTCTTGCA -450
GACAAACTGCGCAACTCGTGAAAGGTAG

**Figure S1. DNA sequences and propensity to bind IHF.**

(A) Primary score from the PSSM scanning analysis (in blue bars) for all matches with double or more probability to find the sequence into the IHF PSSM compared to *E. coli* genome (see Materials and Methods). (B) Complete sequences for the four DNA constructs studied here indicating the sequence for the known IHF specific recognition sites (in yellow), the part considered at the PSSM (underlined text) and the most conserved positions (in bold).

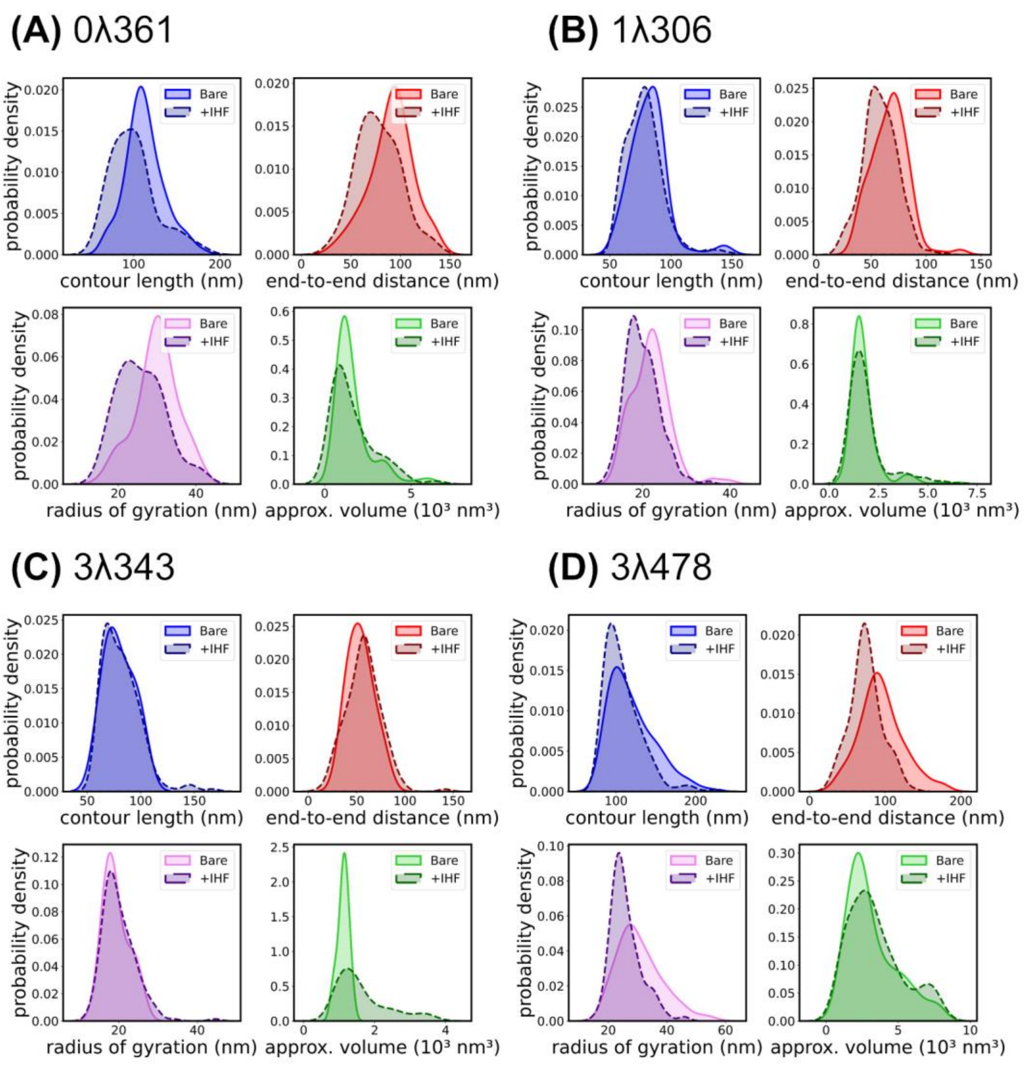


**Figure S2. Additional experimental AFM data.**

Kernel density estimates for dimensional parameters of all DNA constructs used (**(A)** 0λ361, **(B)** 1λ306, **(C)** 3λ343 and **(D)** 3λ478) with and without IHF. Contour length, radius of gyration and end-to-end distances of 1λ306 and 0λ361 present a certain reduction (albeit not statistically significant) upon addition of IHF due to specific and non-specific binding, respectively. For the fragments with three binding sites we observe a combination of several effects on the different parameters. In the absence of IHF, contour length, radius of gyration and end-to-end distances show that 3λ478 is more flexible than 3λ343 as it is longer in terms of DNA persistence length (3.2 vs 2.3 times long approx.), so it behaves more accordingly to the semiflexible regime instead of the rigid regime typical of short DNAs (1). However, in the presence of IHF, these parameters show that DNA is rigidified and that the two constructs tend to converge in the narrow-peaked distribution characteristic of the naked 3λ343. The reduction of DNA flexibility caused by protein binding has already been observed in other protein-DNA complexes like nucleosomes (2). Volume measurements indicate the bigger propensity to cluster of 3λ343 compared with 3λ378 due probably to the elimination of spare DNA that could reduce electrostatic repulsion and steric extrusion between DNA duplexes.

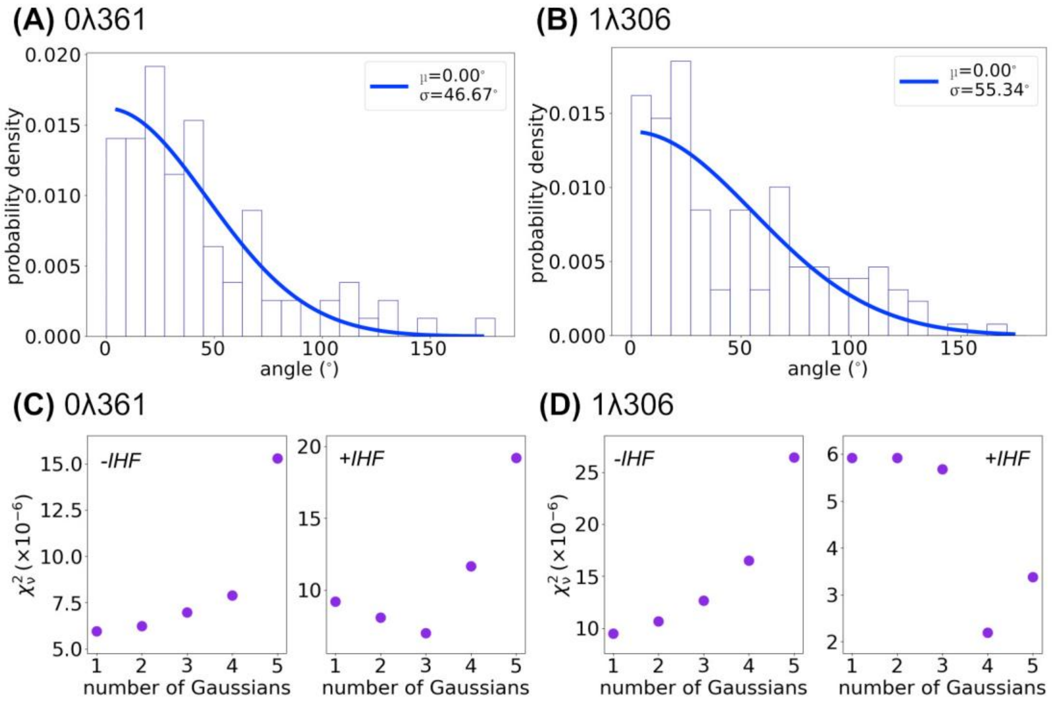


**Figure S3. Additional bending angle analysis**

Angle distributions for **(A)** 0λ361 (*n*=63) and **(B)** 1λ306 (*n*=584) without IHF, showing a Gaussian centered around zero with a width of 47˚ and 55˚ respectively, suggesting that the 1λ306 construct is slightly more flexible, perhaps due to sequence-specific curvature. Reduced $\chi^2$ (one-sided) goodness-of-fit tests for 0λ361 **(C)** and 1λ306 **(D)** +/- IHF, $p > 0.99$. For 0λ361 with IHF, the data is fit well by either 2 or 3 Gaussians, suggesting an unbound and 1-2 bound states. For 1λ306 with IHF, the reduced $\chi^2$ is lowest for four Gaussians, suggesting that there are four states, the fixed unbound state and three bound states. In both cases, without IHF, the data is best fitted by a single gaussian function as shown in (A) and (B).

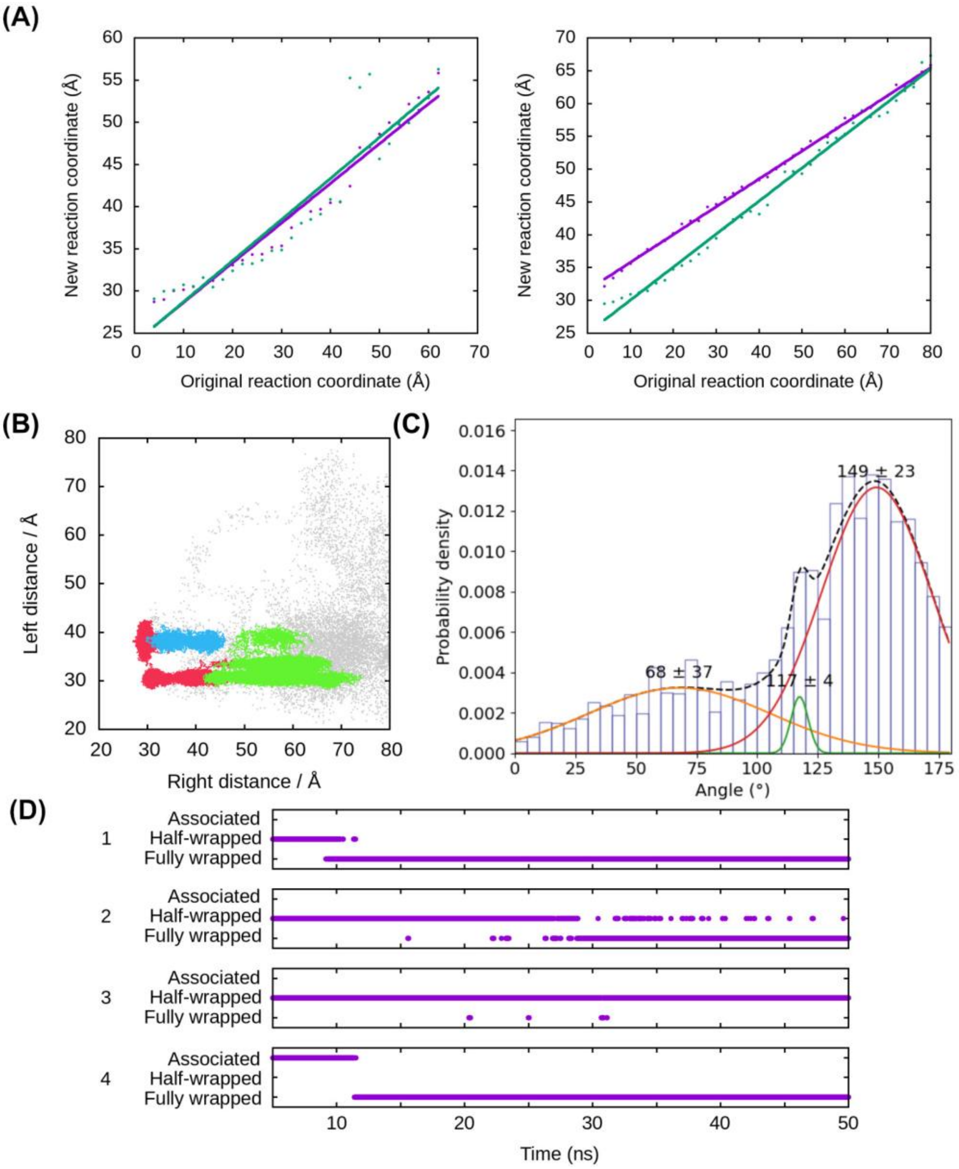


**Figure S4. Additional simulation data.**

**(A)** The original reaction distance chosen for the calculation of PMFs (as described in Materials and Methods) was converted into a new reaction distance between the centers-of-mass of the protein and a section of DNA to eliminate distortions caused by local structural fluctuations. The scale factor between the two reaction coordinates is approximately linear ($R^2$ = 0.84–0.99) on both the left (left) and right (right) sides, whether the other arm is restrained away from the protein (purple) or allowed to move freely (green). The new reaction coordinate is the one used on all free energy plots. **(B)** By identifying the cluster to which each frame of the simulation belongs to**,** the areas representative of each state are delineated onto the free-energy landscape (fully-wrapped in red, half-wrapped in green, associated in blue, snapshots from the first 5 ns of each replica in grey)**. (C)** The bend angle distribution calculated considering the four replicas presents 3 peaks that when they are fitted by gaussian functions are in good alignment with AFM and MD structural clustering. **(D)** The possible transitions can be observed, with every replica spending some time in the associated or half-wrapped state before transitioning to the canonical fully wrapped state.

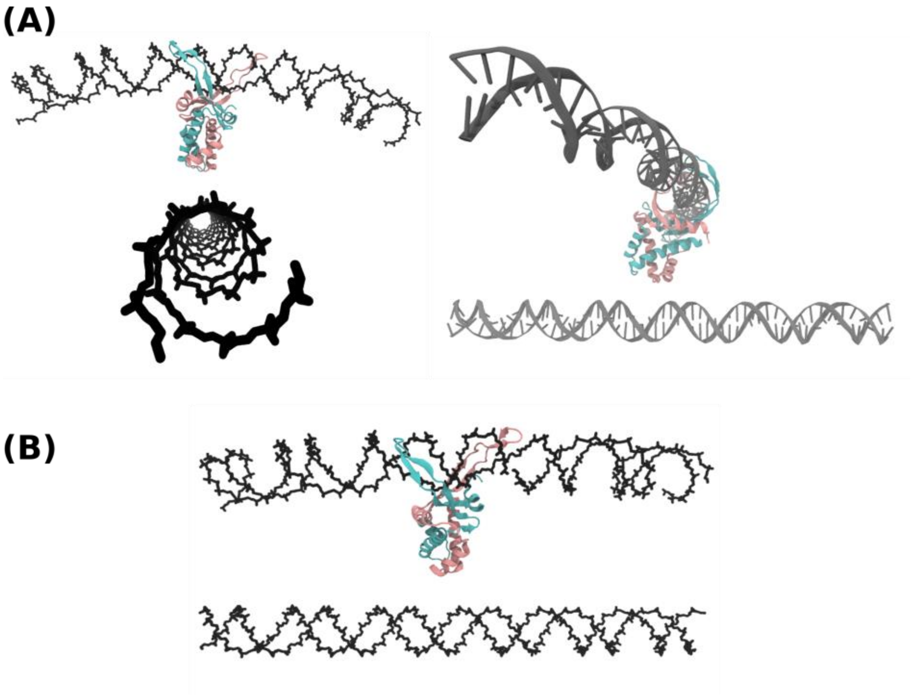


**Figure S5. Initial structures for simulating DNA bridging mediated by IHF.**
**(A)** Two views of the structure used as a starting point for modeling bridge formation where the two 61 bp pieces of double-stranded DNA were perpendicular to each other in order to minimize electrostatic repulsion. **(B)** An example of an additional attempt of initial structure where the two DNA molecules were placed in parallel. This orientation did not produced realistic bridging without allowing major rearrangement.

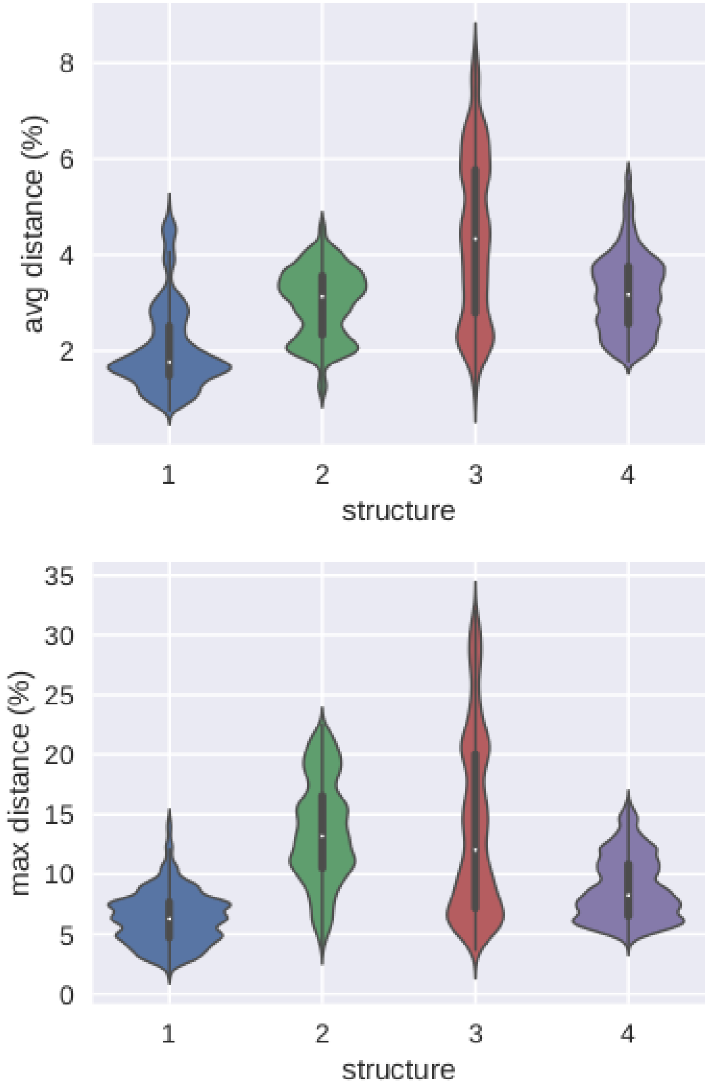


**Figure S6. Deviation from planarity.**
The average (top) and maximum (bottom) distance along the DNA to the best fitted plane is less than 10% and 35%, respectively, in relation to the molecule's height at any frame of all replicas run in implicit solvent of the 1λ306 construct with IHF.

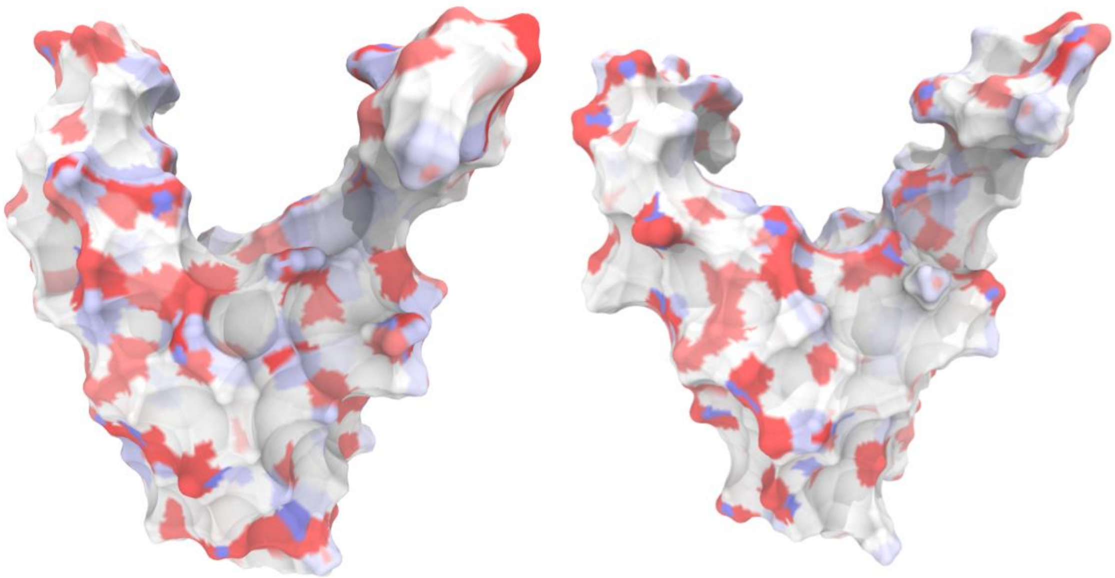

**Figure S7. Surface electrostatic profiles of IHF and HU.**
The surfaces of IHF (left) and HU (right) are similar from an electrostatic perspective, both consisting of isolated spots of positive charge (blue) on an otherwise neutral (white) or negatively charged (red) surface.

### Supplementary Movies

**Supplementary Movie 1. Replica 1: DNA rapidly achieves the fully wrapped state.** Movie of the first replica where the complex rapidly reaches the fully wrapped state (Figure S4D). Because this is the replica with a most canonical behavior, it is selected in Figure 1 for assessing the recovery of crystallographic interactions by means of implicitly solvated simulations.

**Supplementary Movie 2. Replica 2: DNA fluctuates between the fully and half-wrapped states when bound to IHF.** Movie showing the second replica of implicitly solvated simulations of the 1λ306+IHF experimental construct where the complex oscillates between the half- and fully wrapped states (Figure S4D).

**Supplementary Movie 3. Replica 3: The half-wrapped state is a metastable state.** Movie of the third replica where the complex remains in the half-wrapped conformation showing the metastable nature of this state (Figure S4D).

**Supplementary Movie 4. Replica 4: The transition to the fully wrapped through the associated state.** Movie of the fourth replica where the complex traverses through the associated state before transitioning to the fully wrapped state (Figure S4D).

**Supplementary Movie 5. DNA mechanical switch based on dynamic allostery.** Representative movie based on umbrella sampling trajectories of the asymmetric allosteric switch between arms, where the left arm (with the A-tract, in red) binds before the right arm containing the specific recognition sequence (in blue)
